# Quinacrine binds to the kinase domain of FGFR1 and inhibits its activity

**DOI:** 10.1101/2021.12.02.470934

**Authors:** Makhan Kumar, Angshuman Sarkar

**Affiliations:** CMBL, Department of Biological Sciences, Birla Institute of Technology and Sciences, K K Birla Goa Campus, Sancoale, South Goa – 403726, India

**Keywords:** Quinacrine, FGFR1, Proliferation, Migration, Angiogenesis

## Abstract

**Background:** FGF family receptors, especially FGFR1, have been widely implicated for their potential role in the promotion of oncogenesis and chemoresistance in lung cancer. Quinacrine, an anti-malarial drug, has been widely reported to exhibit anti-neoplastic properties through the activation of p53 and simultaneous inhibition of NF-kB signaling pathways in cancer cells.

**Methods:** The binding of QC to FGFR1 was studied using molecular docking and molecular dynamics simulation studies. The experimental kinase activity assay for the protein was performed using a luminescence-based kinase assay. FGF-induced phosphorylation and proliferation were studied by cell counting and western blotting. Matrigel-based cell migration was conducted to assess migration activity.

**Results:** QC interacted with multiple residues around the kinase insert domain of FGFR1 through hydrogen bonding, hydrophobic interactions, and water bridges. The kinase activity inhibition assay demonstrated a significant reduction in FGFR1 kinase activity by QC at higher concentrations, which was further observed at the cellular level in inhibition of FGFR1 phosphorylation and proliferation by QC at higher exposure concentrations of FGF stimulated cells. These effects were further validated downstream in the FGF-induced activation of cell migration.

**Conclusion:** QC did form a stable interaction with residues of FGFR1 at another allosteric site surrounding the kinase domain, leading to inhibition of its kinase activity at higher drug concentrations. This effect was further observed at the cellular level in both acidic and basic FGF ligand-induced proliferation, phosphorylation of FGFR1, and cell migration, where a trend of significant reduction in activity was observed at higher drug concentrations.

## 1. Introduction

The fibroblast growth factor (FGF) family of extracellular growth factors consists of 18 ligands that exert their function by binding to one or all of the four receptor tyrosine kinases (RTK) FGF receptor 1, FGFR2, FGFR3, and FGFR4 [1]. These RTKs are highly conserved, and upon binding with the extracellular ligand, initiate a series of activation cascades that regulate numerous cellular processes such as proliferation, migration, and wound healing. FGF and its receptors play a critical role in embryonic organogenesis, injury repair, and endocrine homeostasis in adults [2]. Upon binding of the FGF ligand to its specific receptor, dimerization of the receptor takes place, leading to initiation of receptor activation via a precise sequence of autophosphorylation. It starts with phosphorylation at the Y653 residue, followed by phosphorylation of Y583 and 585(kinase insert region),463(juxtamembrane region), 766(C-terminal region), and finally Y654 in the activation loop, which facilitates the binding of numerous downstream signaling molecules through Src homology 2(SH2) or phosphotyrosine kinase(PTB) binding domains [3]. FGF1 (acidic FGF) and FGF2 (basic FGF), out of the 18 members of the FGF family, are known to have predominant activity regulating various cellular behaviors, including proliferation, cell cycle regulation, and migration. FGF1 is the only member of the entire family that can bind and activate all four FGFRs, whereas FGF2 has a selective affinity for FGFR1 and FGFR3 receptors [4].

Over the decades of research work since the discovery of FGFs in the 1970s, a large number of research reports have accumulated suggesting aberrations in the FGF-FGFR signaling cascade and their direct implications in oncogenesis, metastatic spread, and acquiring drug resistance. Overproduction of FGFs, particularly FGF2 and FGF1, has been reported to play a role in increased tumorigenicity, EMT, invasion, and metastasis in multiple malignant tumors including nsclc, breast, prostate, and head and neck cancer [5]. Molecular aberrations, such as gene amplification and fusion, have been reported in various cancers, including lung cancer, and are directly associated with aggressive cancer growth and spread [6, 7]. FGFR1, a member of the FGFR family, has been well reported and documented in various malignancies, especially lung cancer. FGFR1 alterations, including gene amplification, were found to be more common than other members of the FGFR family in a cancer genome sequencing study of over four thousand patients [8]. FGFR1 is known to interact and cooperate with EGFR to accelerate tumor growth, and it has also been reported to facilitate resistance by providing an escape from clinically approved EGFR inhibitors such as afatinib in lung cancer [9]. A few other studies have also highlighted the aberrant expression of FGFR1 in response to EGFR and MEK inhibitors, leading to increased EMT and bypass growth pathways via activation of STAT and Akt downstream signaling pathways and subsequent resistance to targeted tyrosine kinase inhibitors [10]. All these preclinical and clinical findings make the FGFR family, especially FGFR1, in the case of lung cancer, an ideal candidate for targeted drug development. Several inhibitors targeting FGFR kinase activity have been developed, and currently, four of them, lenvatinib (for iodine-refractory thyroid carcinoma), ponatinib (drug-resistant CML), pazopanib (renal cell carcinoma and sarcoma), and regorafenib (for colorectal carcinoma and gastrointestinal stromal tumors) have already been approved for clinical use, and several other targeted inhibitors are currently under clinical trials [11].

Quinacrine, a synthetic acridine-based anti-malarial drug, has been well studied and documented for its antineoplastic activity in numerous preclinical and clinical studies [12–14]. The molecular mechanisms of its anti-cancer effects have been reported to be mediated through the activation of p53 signaling, induction of autophagy, DNA and mitochondrial damage, and facilitation of interaction between TRAIL and death receptor DR5 [15–18]. However, these mechanisms do not provide a complete picture of QC antineoplastic activity, as the effect of QC on the crucial aspects of cancer metastasis and chemoresistance is not clearly understood. Apart from phospholipase A_2_, other molecular targets of QC that could have a potential impact on tumorigenicity have not been covered as well. Recently we worked on this aspect and discovered a novel interaction of QC with GSTA1 which encodes the GSTα protein and is well known for its prominent role in cancer cell proliferation, survival, and most importantly chemoresistance [19]. Here, we report a novel interaction of quinacrine with FGFR receptor family member FGFR1 and inhibition of its kinase activity and downstream cellular processes by QC.

## 2. Materials and Methods

### 2.1 Cell culture and preparation of Quinacrine solution

Dulbecco’s modified Eagle medium (DMEM) and fetal bovine serum (South American origin) were procured from HIMEDIA, India. The NSCLC cell line A549 was procured from the National Center for Cell Science (NCCS), Pune, India, and was authenticated by the aforementioned center using the AmpFISTR identifier plus PCR amplification kit from Applied Biosystems, and the percentage match between the tested sample and STR profile database was found to be 95%. The cells were cultured in DMEM supplemented with 10% fetal bovine serum (FBS). The cells were maintained at 37°C with 5% CO_2_ and 95% humidity in a CO_2_ incubator.

Quinacrine dihydrochloride powder purchased from Sigma (Q 3251) was weighed and dissolved in sterile tissue culture grade water to make a 1mM stock solution and was stored at 4 °C under light-protected conditions. Cells were exposed to 5, 10, and 15 μM concentrations per ml of media for all experiments mentioned in this article from this stock solution.

### 2.2 Molecular docking studies of Quinacrine with FGFR1

The crystal structure of FGFR1 (kinase domain) (PDB ID: 4uwy) was downloaded from the RCSB Protein Data Bank [20]. The initial 3D coordinates of the QC were downloaded from the RCSB database. FGFR1 in complex with QC was modeled using the iGEMDOCK molecular docking program [21]. A stable docking option with population size N=300, generations (80), and no. of solution (10) was selected. Repeated studies were performed using the same compound and settings to avoid false-positive and false-negative results.

Post-docking analysis: Modeled protein complexes were studied for a detailed understanding of protein-ligand interactions using PyMOL. PRODIGY-lig, and Protein-Ligand interaction Profiler (PLIP) servers were used to predict the binding affinities of the modeled FGFR1-QC complex based on structural signatures [22].

### 2.3 Molecular Dynamics studies of QC’s binding with FGFR1

The 3D coordinates of the kinase domain of VEGFR2 and Lyn were retrieved from PDB, as described earlier. Three molecular dynamics simulations were conducted for the protein (FGFR1-Ponatinib, FGFR1-QC, and Apo FGFR1), with a length of 100 ns for each MD. Discrepancies in the experimental structure, such as the absence of hydrogens, missing atoms, missing side chains, and selenomethionines, were corrected using the Protein Preparation Wizard tool of the Maestro Visualization tool (academic version) of Schrodinger Software. The terminal amino acids were capped, and the pH of each condition was optimized at pH 7.0. The resulting optimized protein mimics the physiological state. The proteins were then solvated in an orthorhombic box with a periodic boundary condition of 10 Å from the walls of the simulation box, and pre-calibrated water was added through the TIP3P model. The overall neutrality of the solvated system was managed by adding the requisite number of counterions, *that is*, Na+ or Cl- ions. Once the solvated system was neutralized, a physiological salt concentration of 0.15 M was maintained through the addition of an appropriate amount of Na+ and Cl- ions. The MD simulation tool was used to simulate solvated models using the NPT ensemble algorithm. The shortest iteration time of the simulation was set to 2 fs using the RESPA integrator. The coordinates of the system were recorded in the trajectory after every 100 ps, so that a total of 1000 conformations were recorded in the simulated trajectory. The root mean square deviation/fluctuation and radius of gyration plots were generated using the simulation event analysis module of Desmond. Dihedral angle analysis of inhibitors, ligand interaction maps, and protein interaction bar plots were generated using the simulation interaction diagram tool of Desmond.

### 2.4 FGF induced cell proliferation assay

Recombinant aFGF/FGF-1 and bFGF/FGF-2 were purchased from Thermo Fisher Scientific (13241-103 and PHG0024), and 1μg/ml stock solutions were prepared in serum-free DMEM and used for the experiments. Approximately 5×10^4^ cells were seeded separately into 6-well plates and allowed to grow under standard conditions for 24 h. Cells were then treated with 10 ng/ml of acidic and basic FGF procured from Invitrogen. QC (5, 10, and 15 μM) was added to each well, and the cells were incubated for 24 h in a CO_2_ incubator at 37 °C. After incubation, cells were harvested by trypsinization, centrifuged at 1500 rpm for 5 min, and resuspended in 1 ml DMEM. Total viable cells were counted by the trypan blue dye exclusion method using an improved Nauber hemocytometer. Three independent experiments were performed, and the average total viable cell count was plotted against the exposed drug concentration using Microsoft Excel.

### 2.5 FGFR1 kinase enzyme activity inhibition assay

FGFR1 kinase enzyme system kit was purchased from Promega Inc. (V9321) was used for the reaction. ATP and ADP standards for background, ATP/substrate mix, and kinase solution were prepared according to the manufacturer’s instructions. First, the titration of Lyn kinase was performed through a serial dilution starting from 200 ng in duplicates as per the instruction sheet of the product in a white flat bottom 96 well plate and incubated at 37 °C for 60 min. After incubation, the ADP Glo reagent and kinase detection reagent were added to the wells, incubated for 40 min each, and the plates were read in a Perkin Elmer Victor3 multimode plate reader instrument at 0.5 seconds integration, and three separate readings were taken at intervals of 10 s each. The SB10 values (amount required to generate a signal to background ratio of 10) were calculated, and the kinase solution for the calculated SB10 was prepared as per the instruction sheet. The QC inhibitor solution was prepared by serial dilution starting at 10,000 nM. The kinase reaction was allowed to occur at 37 °C for 60 min, and post-incubation was terminated by adding ADP Glo reagent, incubated with kinase detection reagent for 40 min, and the readings were taken using the same instrument mentioned earlier. The inhibition activity graph was plotted, and the IC50 value for QC was calculated using the online IC50 tool [23].

### 2.6 RNA isolation and Reverse Transcriptase-PCR analysis

Ribozol RNA extraction reagent was purchased from Takara Life Sciences and the Verso cDNA Synthesis Kit (Thermo Scientific). The primers used for RT-PCR analysis were designed and checked for specificity using NCBI Primer-BLAST and were ordered from IDT. Total RNA was isolated directly from a 25 cm2 cell culture flask, and cDNA synthesis was performed as previously reported protocol^37^. Specific primers for each gene were 18SrRNA-Forward-5’-GTAACCCGTTGAACCCCATT-3’. Reverse: 5’-CCATCCAATCGGTAGTAGCG-3’, vimentin - Forward-5’-GACAATGCGTCTCTGGCACGTCTT-3’. Reverse-5’-TCCTCCGCCTCCTGCAGGTTCTT-3’,MMP9-Forward-5’-CAGAGATGCGTGGAGAGT-3’. Reverse: 5’-TCTTCCGAGTAGTTTTGG-3’, and N-Cadherin-Forward-5’-CAACTTGCCAGAAAACTCCAGG-3’. Reverse-5’-ATGAAACCGGGCTATCTGCTC-3’ was used to amplify selected gene targets by RT-PCR. 18S rRNA was used as an internal control, and band densitometry analysis was performed using NIH ImageJ software.

### 2.7 Protein extraction and Western blotting

Phenylmethanesulfonyl chloride (PMSF) and Ponceau S, a practical grade powder, were procured from Sigma Life Sciences. Protease inhibitor cocktail tablets (EDTA-free) were purchased from Roche diagnostics. Tris base and all other routine chemicals were purchased from MP Biomedicals, India. Primary antibodies against β-actin (A5441), phosphor-FGFR1 Tyr 653/654 (06-1433-25 μg), and total FGFR1 (F5421) were purchased from Sigma Life Sciences. The primary antibodies for N-cadherin (13116T) and E-cadherin (3195T) were purchased from Cell Signaling Technology (Danvers, MA, USA). Primary polyclonal antibodies against MMP9 (E-AB-10063) and vimentin (SC-6260) were purchased from Santa Cruz Biotechnology. HRP-linked secondary antibodies (NAV 930 and 931) were purchased from GE Healthcare UK Limited. Total cellular protein was extracted using lysis buffer (Tris 10mM pH7.4, EDTA, 1mM, pH 7.4, PMSF, protease inhibitor cocktail tablet, Triton X-100) on an ice bath, followed by centrifugation at 16,128×g for 20 min at 4°C. Quantification was performed using the Bradford method, and approximately 40μg of cell lysate was run and normalized on a 10 %–12% SDS-PAGE gel. The resolved gel was then transferred to a polyvinylidene fluoride membrane and processed as previously reported [19].

### 2.8 Matrigel cell migration assay

Cells were grown in a 6-well plate, treated with various concentrations of QC, and stimulated with 0 ng FGF, 10 ng aFGF, and 10 ng bFGF for 24 h. Permeable migration chambers purchased from Corning Inc. were coated with 50 μL of 50 μg/ml Matrigel and incubated at 37°C for 24 h. After that, 5×10^4^ cells of each sample and untreated control were suspended in 200μl of media without FBS and added to the top chamber. The migration chambers were then kept in 24 well plates with media and FBS and kept in a CO_2_ incubator for 6 h, allowing cells to migrate. Cells were then fixed with 4% formaldehyde and stained with 2% crystal violet, followed by washing with PBS. Images of the migrated cells were captured on an Olympus inverted microscope at 10× magnification, and migrated cells were counted from five different fields from each of the samples using NIH ImageJ software.

### 2.9 Statistical analysis

All statistical analyses were performed using GraphPad Prism 6.0 (Graph Pad Software Inc. La Jolla, USA). Values are expressed as the mean ± standard deviation (SD). Data were analyzed using ANOVA, followed by multiple comparisons that were carried out using the Bonferroni post-hoc test. Statistical significance was set at p < 0.05.

## 3. Results

### 3.1. Quinacrine interacts with residues surrounding kinase insert domain of FGFR1 in molecular simulation models

Quinacrine interacts with one of the FGFR family receptors, FGFR1. The binding energy values obtained for the FGFR1-QC complex were calculated to be −72.4 kcal/mol. The PRODIGY-Lig server predicted the binding affinity of the modeled FGFR1-QC to be 7.8. Further, a detailed analysis of residue interactions revealed that residues Tyr572, Glu593, Gln594, Leu595, and Asn 635 of FGFR1 formed hydrophobic (π-alkyl and alkyl) interactions with the C, N, and O atoms of the planar rings of quinacrine (Supplementary Table 1). Molecular dynamics simulations for a duration of 100 ns were performed to further understand the nature and stability of the QC-VEGFR2 interaction. QC was found to bind with FGFR1 through two hydrogen bond interactions with residues Asp524 and Asp527, a water bridge through Asp641 and Leu 644, and hydrophobic interactions through Phe489 and Leu516. The RMSD of the QC-FGFR1 complex was found to be within 4 Å throughout the simulation (Figure 1a, 1b and 1c). The relatively larger fluctuations were observed in apo state of kinase compared to the complex with inhibitor, suggesting that binding of the inhibitor locks the kinase and restricts its dynamics at activation loop (*i.e* residues 560-590). This region has been known to regulate the catalytic functions and conformational transitions in kinases. The dihedral angle, which is used in MD simulation to identify freely rotatable groups, was found to be connected to terminal aliphatic groups; however, they were seen to be freely rotatable because they have lower steric constraints. The torsion of the drug (QC) was observed to be concentrated around a certain value, suggesting that both drugs are accommodated well in their binding pockets (Figure 2a, 2b, 2c, and 2d). To confirm these computational interactions at an experimental level, an FGFR1 kinase activity inhibition assay was performed to experimentally validate the findings of the molecular docking results of QC interaction with the TK domain of FGFR1. The luminescence activity corresponding to total ADP produced by the kinase was seen to marginally decrease with the increasing concentration of the drug until 1000 nM, beyond which it showed a sharp reduction until the final concentration. The IC_50_ value estimated for the kinase activity inhibition was found to be 1053.58nM [Figure 3a].

**Figure 1.**
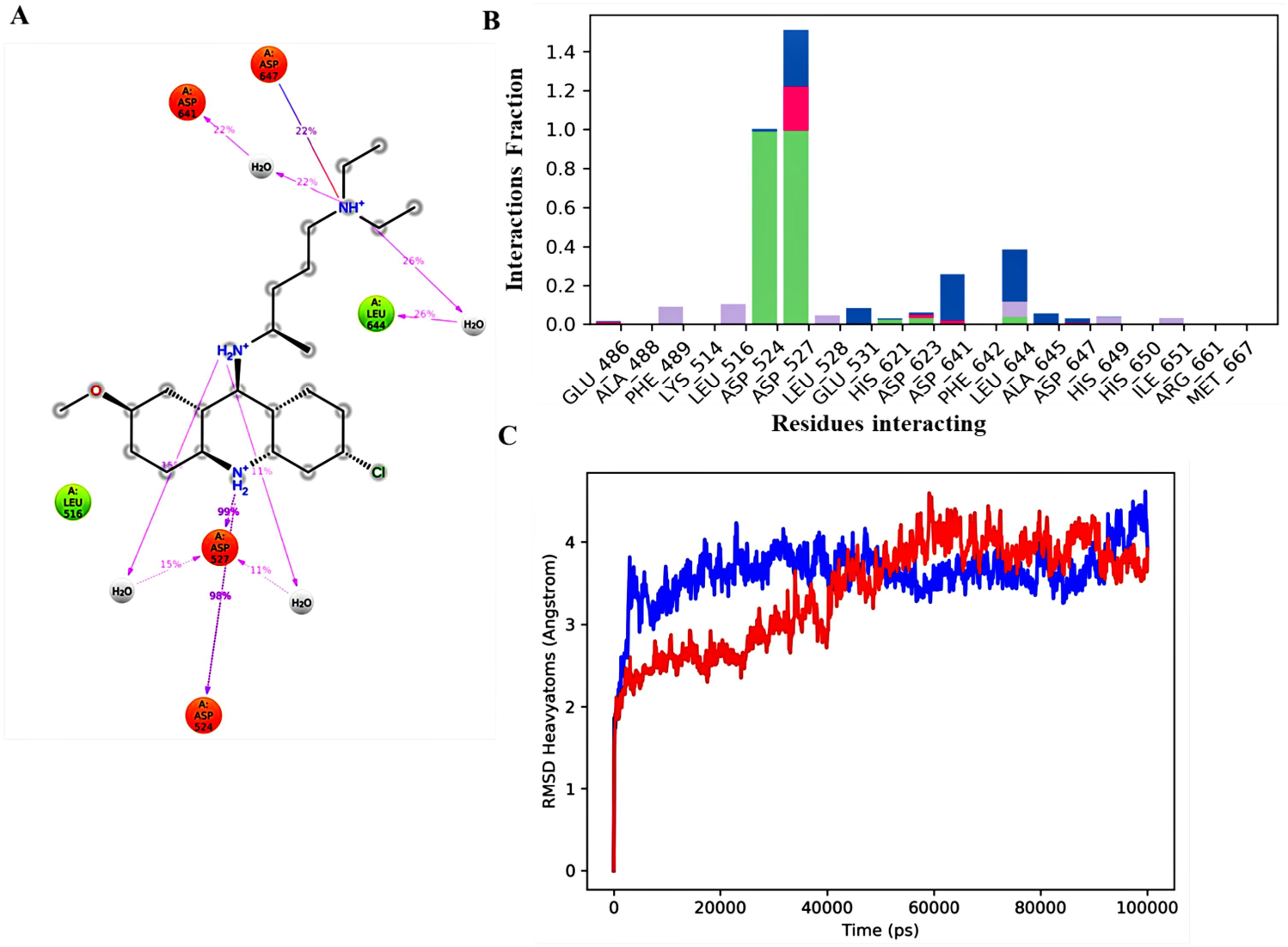
(a) Ligand interaction diagram of quinacrine with FGFR1 kinase domain. Here the amino acids are characterized as negatively charged (rusty red),hydrophobic (lime), polar (cerulean), H-bonding is shown is purple and pi-pi stacking is shown with green lines. (b) Diagram representing the bar plot of physical contacts of drug QC with residues of FGFR1 kinase domain throughout the 100ns long MD trajectory. (c) Root mean square deviation plot for all MD simulations i.e FGFR1 kinase in apo state (shown in red), complexed with and complex with quinacrine (shown in blue) The RMSD trajectories in all the MDs converged around 20 nanoseconds (ns).

**Figure 2.**
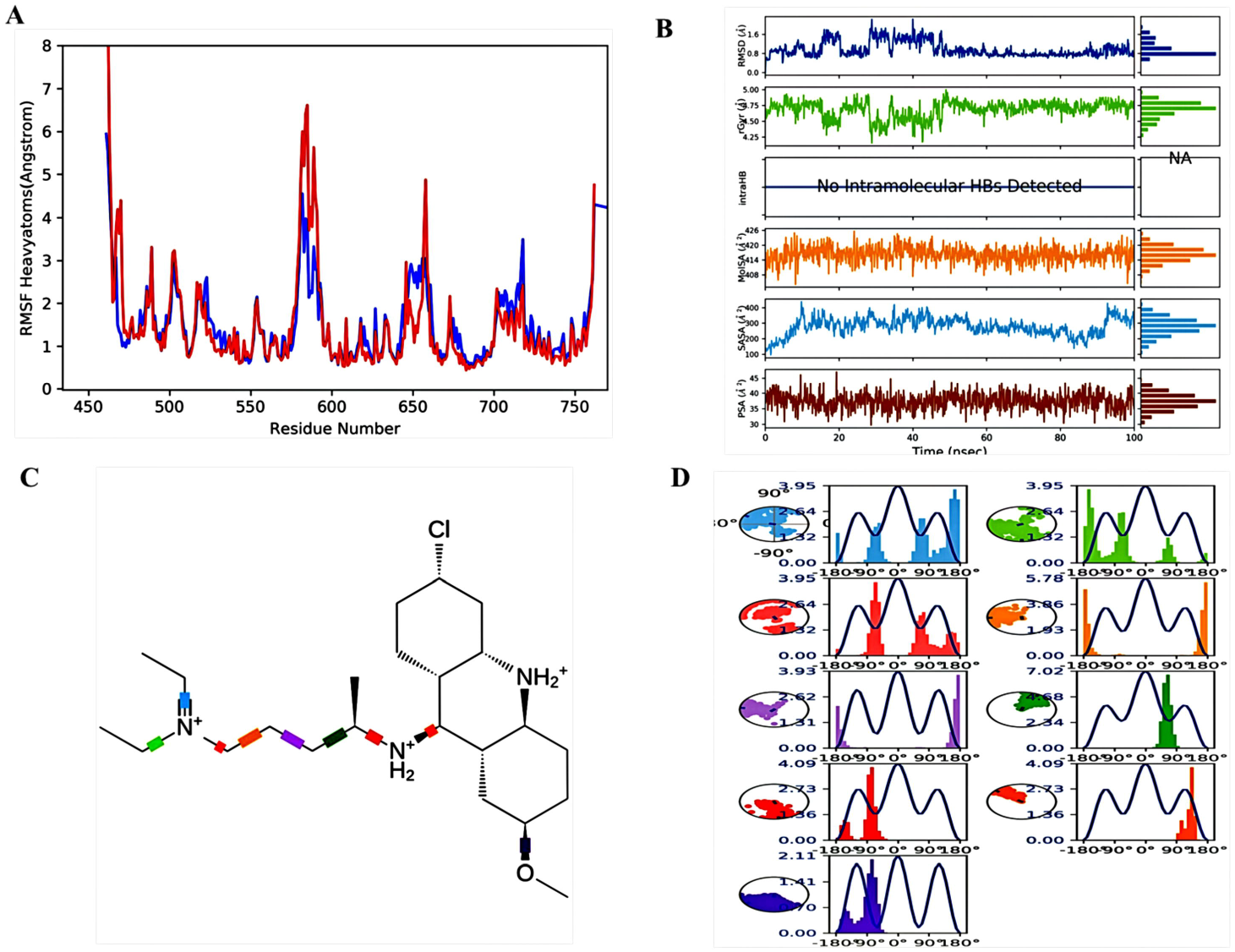
(a) Root mean square fluctuation analysis for all MD simulations i.e FGFR1 kinase in apo state (shown in red and in complex with quinacrine (shown in blue). (b) From top to bottom, the ligand’s root mean square deviation, radius of gyration, intramolecular hydrogen bonds, molecular surface area, solvent accessible surface area and polar surface area are shown. (c) and (d) Dihedral angle plots for QC-FGFR1 complex MD simulations.

**Figure 3.**
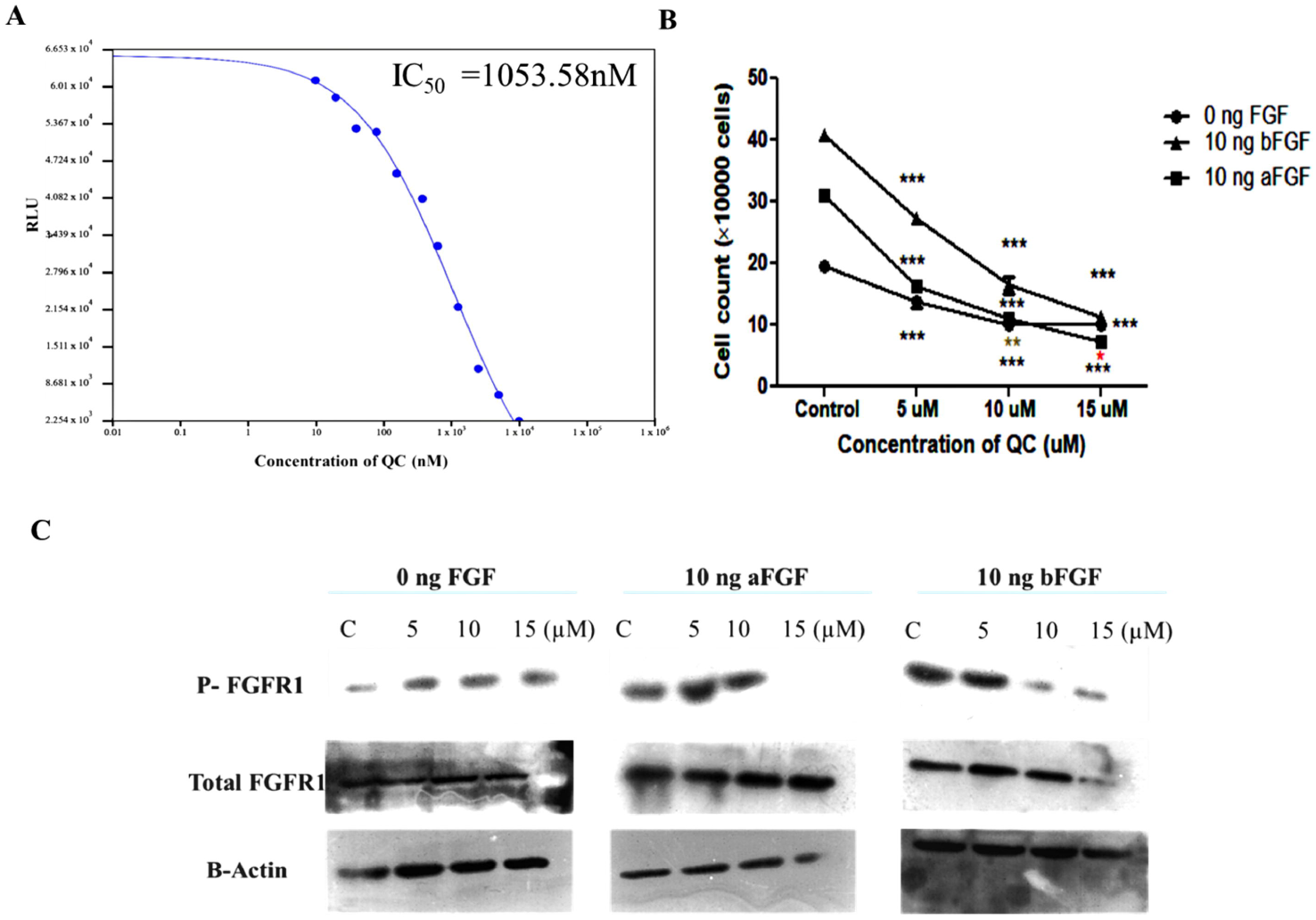
(a) Graph representing luminescence-based kinase activity inhibition of the FGFR1 protein by quinacrine. (b) Graph representing the aFGF and bFGF induced proliferation assay with exposure to different concentrations of quinacrine. Cells were seeded in 6-well plate and treated with various concentrations of QC as described in materials and methods section for 24 h. *=P value<0.05, **=P value<0.001, ***=P value<0.0001 versus control group. (c) Image panel showing western blots of total and phoshopylated FGFR1 in QC treated cells stimulated with 10ng/ml of aFGF and bFGF.

### 3.2 Quinacrine inhibits FGF induced proliferation and phosphorylation of FGFR1

To further confirm the kinase activity inhibition of FGFR1 by QC on the cellular level, FGF induced phosphorylation of FGFR1 and FGF induced proliferation of A549 cells in presence of various concentrations of QC were studied to analyze the QC mediated activity inhibition of FGFR1. Quinacrine significantly inhibited the FGF-induced proliferation of A549 cells, as observed in cells stimulated with 10 ng/mL aFGF and 10 ng/mL bFGF (Figure 3b). A significant reduction of approximately 50% viable cells was observed in cells exposed to 10 μM QC through both FGF-stimulated samples compared to their respective controls. bFGF showed escalating proliferation at faster rates compared to aFGF; however, the effect of QC throughout all doses remained similar across FGF untreated and both a and b FGF-treated samples. Furthermore, the phosphorylation status of FGFR1 (Y653/654), which is a direct consequence of FGF-FGFR1 binding, was studied in QC-exposed cells stimulated with aFGF and bFGF ligands. In aFGF-stimulated and FGF-untreated cells, however, an increase in the quantity of p-FGFR1 was observed at a concentration of 5 μM, which was reduced in subsequent exposure concentrations. In bFGF-stimulated cells, a distinct pattern of dosedependent reduction of p-FGFR1 was observed in QC-treated cell lysates (Figure 3c and Supplementary figure S1).

### 3.3 Quinacrine inhibits FGF induced cell migration and angiogenesis

Activation of EMT and invasiveness of malignant cells is one of the key aspects of FGF-induced activation of FGFR receptor signaling pathways. To understand whether QC-mediated inhibition of FGFR1 kinase activity could affect this cellular phenomenon, matrigel-based cell migration and angiogenesis assays were performed to study this effect. QC was found to inhibit the cell migration process in a dose-dependent manner in FGF-untreated and FGF-treated samples in a manner that was relatively similar across FGF-stimulated and non-stimulated cell samples. The initial exposure concentration of 5μM was observed to induce nearly two-fold reduction in the no. of migrated cells, which further increased beyond a threefold reduction in the 10 μM exposure concentration. The reduction patterns were statistically significant across all doses of QC compared to the untreated control throughout the FGF treatments (untreated, 10ng aFGF treated, and 10ng bFGF treated) (Figure 4a and 4b). To further understand this inhibition mechanism at the molecular level, expression profiles of migration-promoting genes vimentin, N-cadherin, and MMP9 were studied at both the transcriptional and translational levels of gene expression. Significant dose-dependent downregulation of the expression of all three genes was observed in FGF-non-stimulated cells at both the mRNA and protein levels. In contrast, in both aFGF-and bFGF-stimulated cell samples, a marginal downregulation of expression was observed in the 5 μM treatment, which stagnated with increasing concentrations. The expression of vimentin, however, followed a similar trend in non-stimulated cells in both acidic and basic FGF-treated samples (Figure 4c, 4d, Supplementary figure S2, and S3).

**Figure 4.**
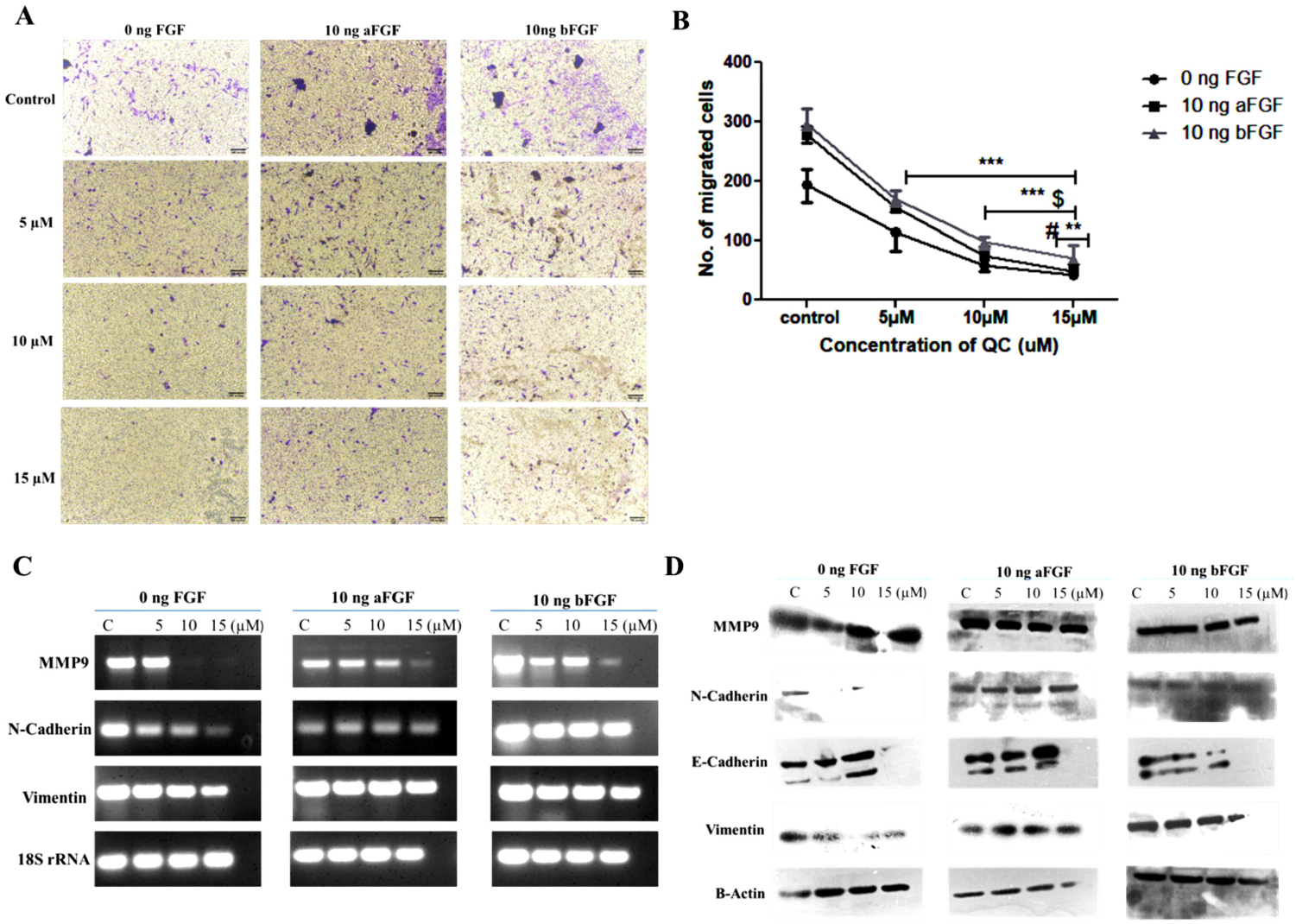
(a) Analysis of QC’s effect on cell migration capabilities of A549 cells stimulated with 10ng/ml of aFGF and bFGF, by matrigel cell migration assay. (b) Graphical representation of no. of migrated cells in QC treated and untreated samples of A549 cell line. The data presented are the mean (±SD) of three independent experiments. (c) Analysis of QC’s effect on mRNA level expression of migration regulatory genes Vimentin, E&N-cadherin for 24 h time point by RT-PCR. (d) Protein level expression analysis of migration regulatory genes after QC exposure for 24 h by western. *=P value<0.05, **=P value<0.001, ***=P value<0.0001 versus control group,$=5 μM versus rest of treatment group #= 10 μM versus rest of treatment group.

## 4. Discussion

The FGF receptor family is well known for its crucial role in promoting cancer proliferation, migration, and chemoresistance. FGFR1, a member of the FGF family of receptors, has been extensively linked to multiple hallmarks of cancer in various cancer types, including nsclc, and has been shown to play a critical role in the prognosis of early stage NSCLC [24]. FGFR1 signaling has been shown to interact with the Hippo/YAP1 pathway, promoting stemness of cancer stem cells, and is also known to regulate Cyclin D1 mediated proliferation and migration of lung cancer cells [25,26]. FGF2-FGFR1 autocrine loop activation has been shown to reactivate ERK and Akt signaling, facilitating resistance to targeted therapies in lung cancer [27,28].

Our results indicate that quinacrine interacts with the residues surrounding the kinase insert domain, which could interfere with the activation loop and subsequently the kinase activity of FGFR1. This possibility was tested experimentally via the kinase activity inhibition assay, which demonstrated QC-mediated inhibition of the kinase activity of FGFR1 at significantly higher concentrations with an IC_50_ value of 1053.58 nM. A higher concentration of QC required to significantly inhibit kinase activity can be expected, as QC was observed to form only hydrophobic and slat bridge-mediated interactions with residues, and the binding site of QC was seen slightly away from the activation loop residues to an allosteric site. This effect of inhibition at relatively higher concentrations was established at the cellular level via the inhibition of proliferation and FGFR1 phosphorylation at Y653/654 residues in acidic and basic FGF growth factor-stimulated nsclc cells. QC was shown to significantly inhibit the proliferation of both a and bFGF stimulated cells at 10 and 15 μM exposure concentrations compared to their respective untreated controls, and similar patterns of inhibition of FGFR1 phosphorylation were also observed in both acidic and basic FGF stimulated cells. Furthermore, the effect of FGFR1 activity inhibition by QC was also observed, translating into the cell migration phenomenon of FGF-stimulated cells, where QC at higher concentrations was observed to significantly inhibit the no. of migrated cells in FGF-stimulated samples in a pattern similar to that of FGF-unstimulated cells. The molecular expression profiles of migration-promoting genes displayed significant dose-dependent downregulation of vimentin in FGF stimulated and unstimulated cells, whereas the expression profiles of N-cadherin and MMP9 showed a very marginal decline in both FGF-stimulated cells in response to QC, suggesting a limited effect of QC on certain genes when treated with growth factors, and the roles of other factors and molecules involved in the inhibition of migration, as seen in the migration assay.

In summary, quinacrine demonstrated the potential to inhibit the kinase activity of FGFR1 at multifold higher concentrations than specific inhibitors of the FGFR family of receptors such as ponatinib, dovitinib, and rogaratinib, which have been shown to achieve inhibition at very low concentrations (IC_50_ values ranging 5-50nM) [29]. However, quinacrine possesses the advantage of targeting multiple signaling pathways and inducing cell death via activation of multiple apoptotic and autophagic signaling cascades, and the scope of inhibiting the activities of specific kinases such as FGFR1 and other enzymes, including GSTα, which we have reported earlier, adds promising value to its expanding horizon of antineoplastic potential, which could be further utilized in designing combinational therapies for better and targeted destruction of cancer with fewer side effects.

## Data Availability and materials

All the research-related experimental data are with authors and have not been submitted or published anywhere else, including preprint websites.

## Author contribution statement

All experiments were planned by MK and AS, and performed by MK. MK wrote the manuscript. AS proofread the manuscript and performed the final editing. Both authors have approved the final version of the manuscript.

## Ethics approval and consent for participation

This study does not involve any animal experiments or human samples; therefore, no ethical clearance was required and taken for the research work.

## Acknowledgments

MK acknowledges UGC JRF [No. 3374/(NET.DEC 2014)] for the fellowship support. The authors also thank Dr. Sukanta Mondal for his guidance in interpreting the computational modeling results.

---

Supplementary Table 1- Table representing the detailed molecular interactions of QC-FGFR1 complex.

Supplementary Figure S1- Graphical representation of the desnitometric analysis of the western blots of QC treated cells stimulated with 10 ng aFGF and bFGF as shown in figure 3(c).

Supplementary Figure S2- Graphical representation of the densitometric analysis of the RT-PCR results as shown in figure 4. The data presented is average of three independent experiments.

Supplementary Figure S3- Graphical representation of the densitometric analysis of the Western results as shown in figure 4. The data presented is average of three independent experiments.

