## Supplementary figures and images for "Quinacrine binds to the kinase domain of FGFR1 and inhibits its activity"

### Supplementary Figure 1

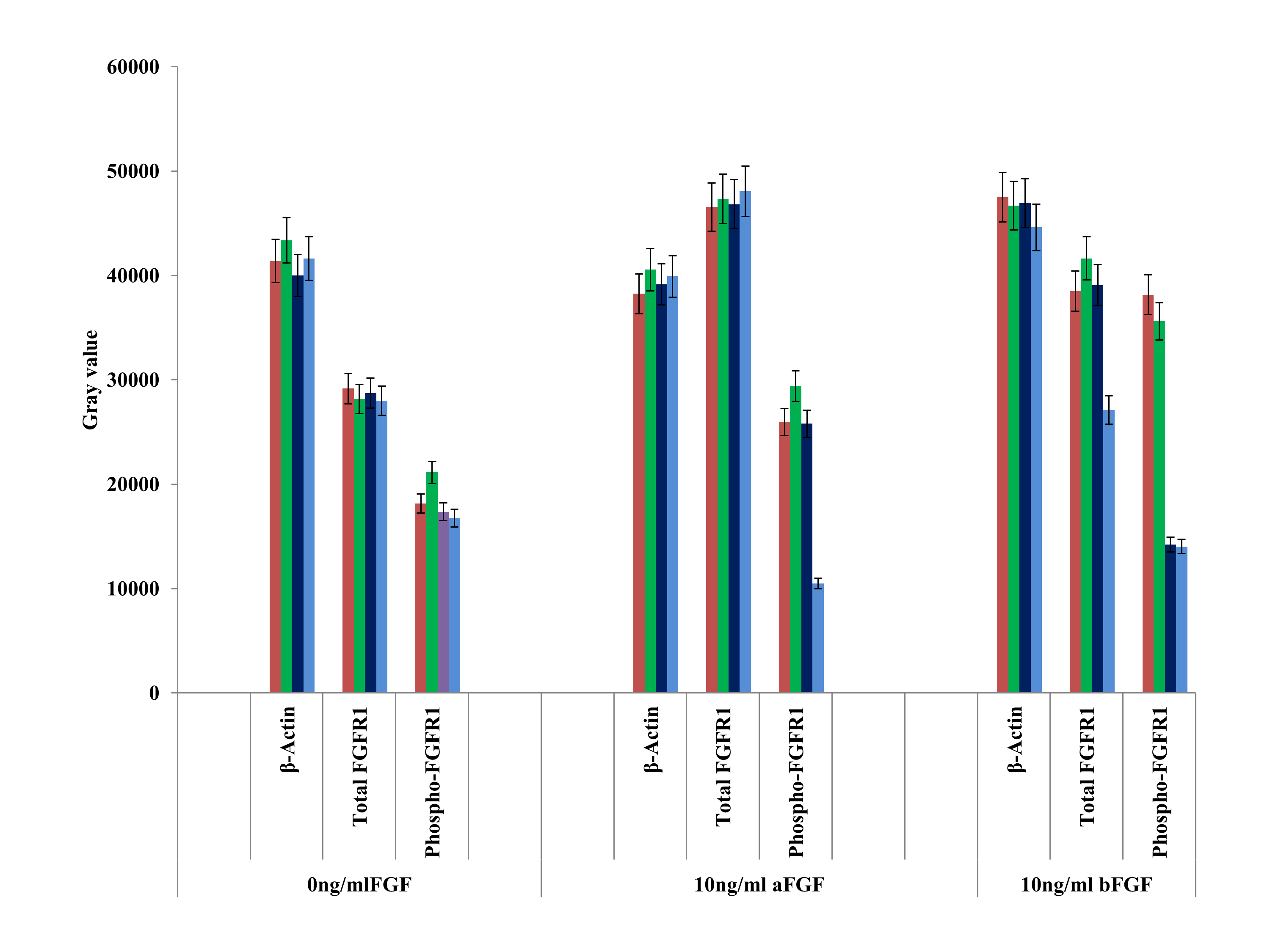

### Supplementary Figure 2

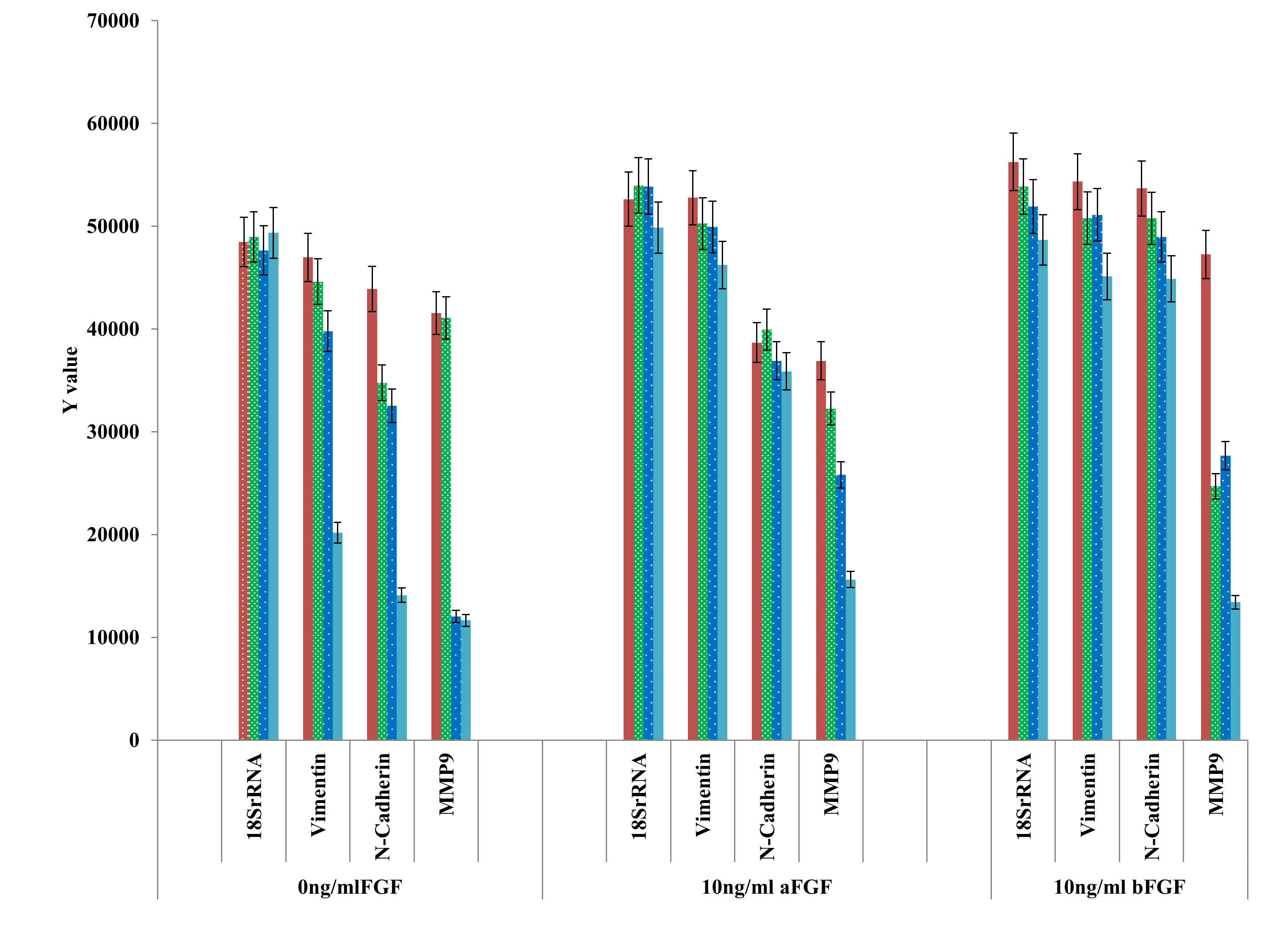

### Supplementary Figure 3

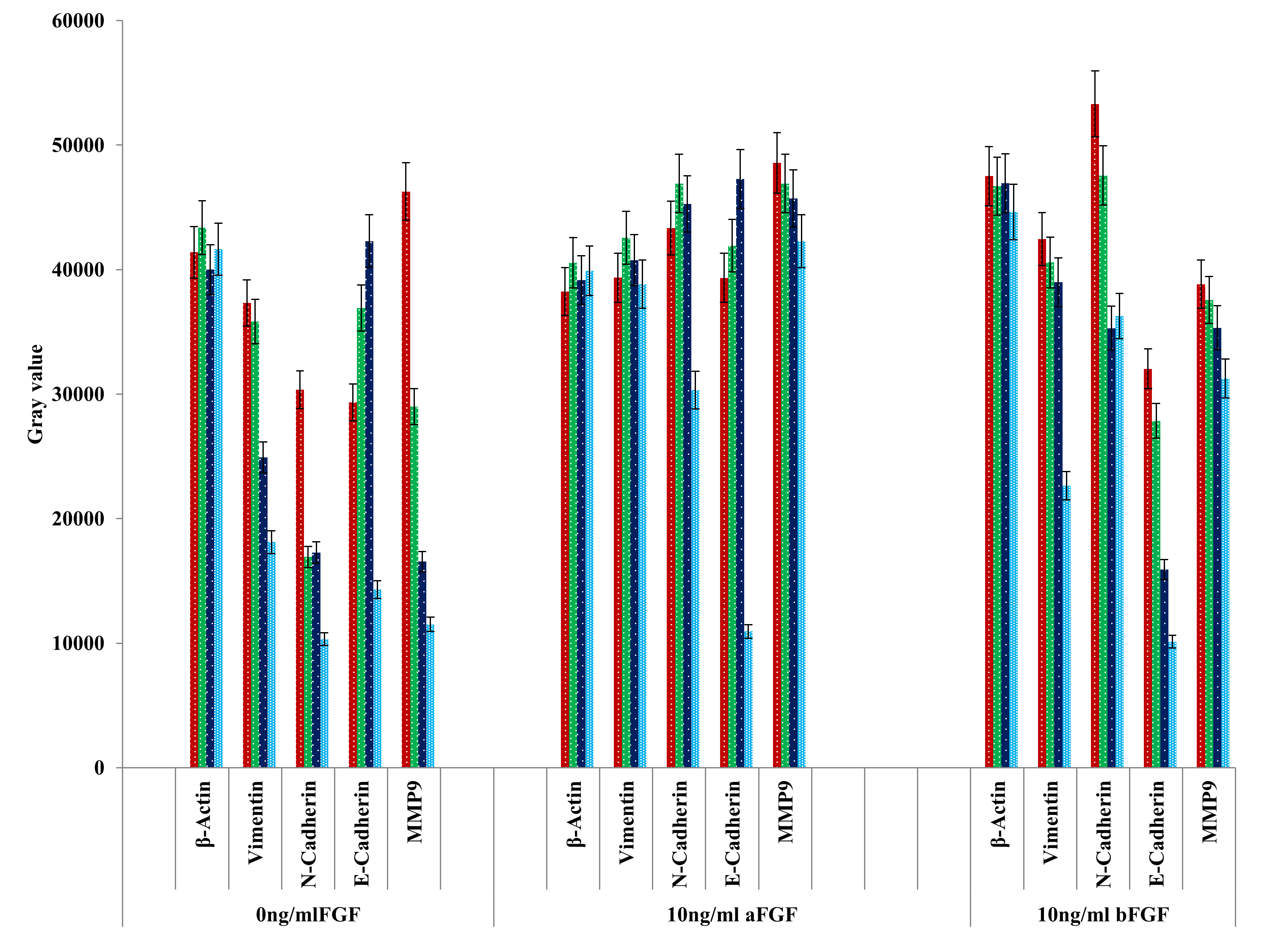
