## Supplementary Table 1 for "Quinacrine binds to the kinase domain of FGFR1 and inhibits its activity"

Table. Modeled molecular interactions of QC, with FGFR1.

| **Bound ligand** | **Predicted affinity(PRODIGY server)**  **Kcal/mol))** | **Interaction** | **Nature of interaction** | **Distance (Å)** |
| --- | --- | --- | --- | --- |
| Quinacrine(QUN) | -7.8 | [A]Tyr572 – QC:N | Hydrophobic (alkyl) | 3.31 |
|  |  | [A]Glu593 – QC: O | Hydrophobic (alkyl) | 3.23 |
|  |  | [A]Gln594 – QC:C | Hydrophobic (alkyl) | 3.40 |
|  |  | [A]Leu595 – QC: O | Hydrophobic (alkyl) | 5.19 |
|  |  | [A] Asp599 – QC:C | Salt Bridges | 3.62 |
|  |  | [A]Ser602 – QC:N | Hydrophobic (π-alkyl) | 6.56 |
|  |  | [A]Asn635 – QC:N | Hydrophobic (alkyl) | 3.73 |
